# Machine learning for discovery: deciphering RNA splicing logic

**DOI:** 10.1101/2022.10.01.510472

**Authors:** Susan E. Liao, Mukund Sudarshan, Oded Regev

## Abstract

Machine learning methods, particularly neural networks trained on large datasets, are transforming how scientists approach scientific discovery and experimental design. However, current state-of-the-art neural networks are limited by their uninterpretability: despite their excellent accuracy, they cannot describe how they arrived at their predictions. Here, using an “interpretable-by-design” approach, we present a neural network model that provides insights into RNA splicing, a fundamental process in the transfer of genomic information into functional biochemical products. Although we designed our model to emphasize interpretability, its predictive accuracy is on par with state-of-the-art models. To demonstrate the model’s interpretability, we introduce a visualization that, for any given exon, allows us to trace and quantify the entire decision process from input sequence to output splicing prediction. Importantly, the model revealed novel components of the splicing logic, which we experimentally validated. This study highlights how interpretable machine learning can advance scientific discovery.

## Introduction

Machine learning algorithms, in particular neural networks, capture complex quantitative relationships between input and output. However, as neural networks are typically black box, it is difficult to extract post-hoc insights on how they achieve their predictive success. Furthermore, they easily capture artifacts or biases in the training data, often fail to generalize beyond the datasets used for training and testing, and do not lead to new insights on the underlying processes^1^.

In recent years, neural networks have been used to tackle challenging biological questions. One outstanding question in genomics is understanding the regulatory logic of RNA splicing, which plays a critical role in the fundamental transfer of information from DNA to functional RNA and protein products. Splicing removes introns and ligates exons together to form mature RNA transcripts. While some canonical sequence features are necessary for exon definition (splice sites delimiting exons and branch points used during intron removal), exon definition is also facilitated by exon sequence^2,3^. Despite recent success using neural networks to predict splicing outcomes^4,5^, understanding how exon sequence dictates inclusion or skipping remains an open challenge. The challenge is further underscored by the sensitivity of splicing logic, where almost all single nucleotide changes along an exon can lead to dramatic changes in splicing outcomes^6,7^.

To enable scientific progress, machine learning models should not only accurately predict outcomes, but also describe how they arrive at their predictions. Here we demonstrate that an “interpretable-by-design” model achieves predictive accuracy without sacrificing interpretability, captures a unifying decision-making logic, and reveals novel splicing features.

### Generating a synthetic dataset for interpretable machine learning

As neural network performance and interpretability is inextricable from the data it is trained on, we began by generating a large, high-quality synthetic splicing dataset. The use of synthetic datasets offers several advantages over genomic data used in previous work. First, genomic datasets are limited by the number of exons in the genome. In contrast, synthetic assays can dramatically increase the number of data points by orders of magnitude^8,9^. Second, genomic exons are flanked by varying sequences (splice sites, introns, promoters) that also participate in splicing decisions^10^, greatly complicating attempts at interpretability. In contrast, synthetic datasets fix all but one variable region, allowing to focus on the region of interest. Third, genomic exons contain overlapping RNA codes (e.g., protein coding sequences). In contrast, sequences in synthetic datasets are devoid of overlapping codes by design. In summary, from both a quantity and quality perspective, synthetic datasets provide crucial advantages for machine learning over genomic datasets.

The synthetic dataset we generated includes hundreds of thousands of input-output data points. Each data point is a different random 70-nucleotide exon sequence, paired with a measured percent spliced in (PSI) output, which is a number between 0 (always skipped) and 1 (always included) (Fig. 1a). The dataset is generated by a massively parallel reporter assay that allows for PSI quantification for hundreds of thousands of unique sequences in a single biological experiment (Fig. 1b). Splicing outcomes for the parallel reporter assay were measured after transfection into human HeLa cells using high-throughput sequencing. We verified that reporters are evenly represented in the reporter assay (Extended Data Fig. 1a). The vast majority of splicing products corresponded to exon inclusion or exon skipping products (Extended Data Fig. 1b), and we filtered our data to exclude spurious splicing products. PSI values are calculated as the number of inclusion reads divided by the total number of inclusion and skipping reads. Three biological replicates of the assay showed excellent agreement (Extended Data Fig. 1c) and their sequencing results were combined for all downstream analysis. High-throughput sequencing measurements were consistent with semi-quantitative measurements of individual reporters (Extended Data Fig. 1d).

**Figure 1.**
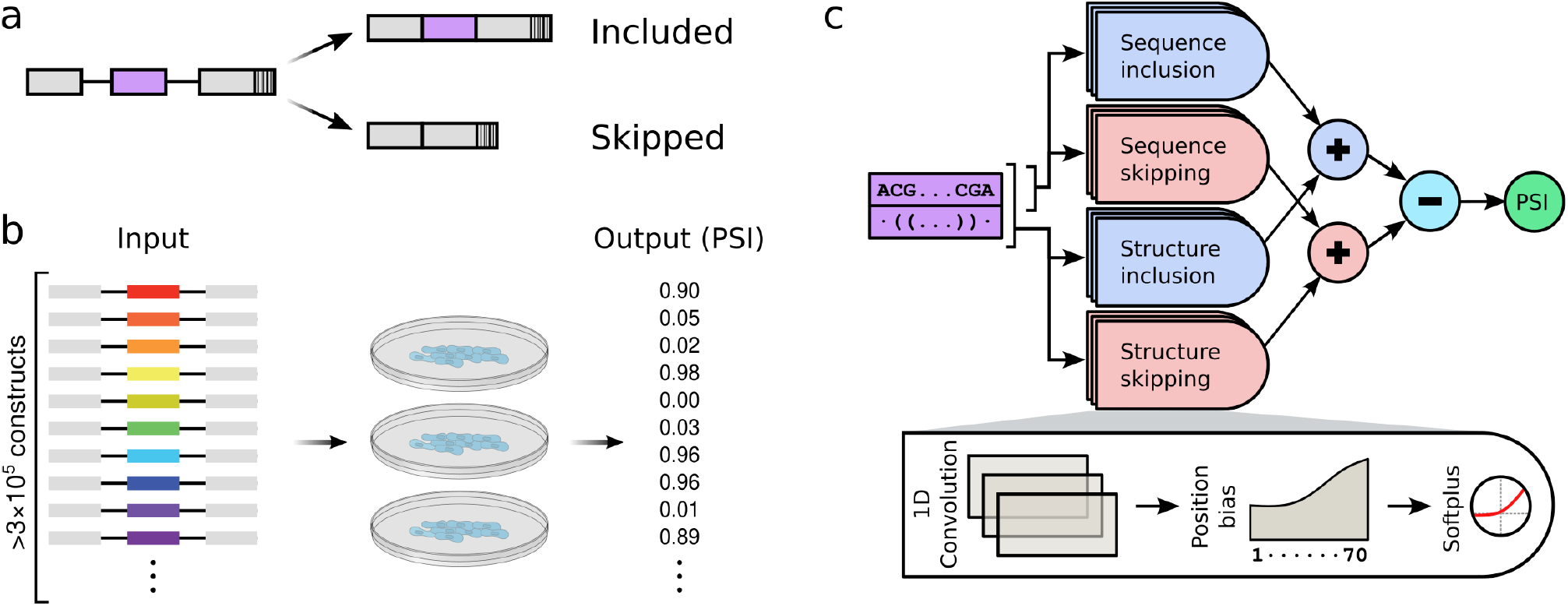
Data generation and interpretable-by-design machine learning model. **a,** All reporters in the assay share the same three-exon design, and differ only in their middle exon, which contains a random 70 nucleotide-long sequence. Depending on its sequence, an exon might be included, skipped, or a probabilistic mix of the two. Each reporter includes a unique barcode at the end of the third exon so that exon identity can be inferred in exon skipping products. **b,** The assay includes over 3 × 10^5^ different reporters. The reporters were transfected into HeLa cells in a pooled fashion in three biological replicates. High-throughput sequencing then provides a “percent spliced in” (PSI) value to each reporter. **c,** The machine learning model consists of both short convolution filters (applied to exon sequence only) and long convolution filters (applied to both exon sequence and predicted structure). The output of these filters (strength) can depend on the position along the exon. Half of the filters are designated as inclusion filters, and the rest are skipping filters. The difference between the total strength of the inclusion filters and the total strength of the skipping filters is used to compute the output predicted PSI.

### An interpretable-by-design model accurately predicts splicing outcomes

State-of-the-art neural networks (based on gated recurrent units^11^ and transformers^12^) trained on this dataset provided excellent prediction accuracy on a held-out test set (RMSE=0.165 and RMSE=0.183, respectively). However, these models are not interpretable, and do not provide any biological insights. We therefore designed a novel model with the explicit goal of being interpretable.

The predictive accuracy of our interpretable-by-design model is comparable to that of state-of-the-art models trained on the same synthetic dataset (RMSE=0.180; Extended Data Fig. 2a). This suggests that interpretability need not come at the expense of accuracy. In addition to our own dataset, the model accurately predicts splicing outcomes from other splicing datasets^7,8,13–16^ (Extended Data Fig. 2b). Importantly, unlike our random exons, these datasets were modeled on specific genomic exons, with each dataset differing in splice sites, introns, and flanking exons. Furthermore, these datasets were generated in different immortalized cell lines. Encouragingly, despite these dramatic differences in RNA architecture and cell types, our model tested well on these datasets, suggesting that our model generalizes and captures critical aspects of splicing regulatory logic.

### Model architecture reveals unifying decision-making process

Our interpretable-by-design model incorporates domain knowledge throughout its architecture (Fig. 1c). Specifically, we reasoned that short six nucleotide sequence filters would capture motifs previously demonstrated to play an important role in splicing decisions^17,18^. We therefore introduced one-dimensional convolutional filters applied to the input RNA sequence. Next, since RNA secondary structure was previously implicated in splicing outcomes^15,19^, we also provided the network with predicted structure^20^. We then introduced longer (30 nucleotide) one-dimensional convolutional filters to the structure-augmented sequence. Crucially, while we fixed filter lengths using minimal domain knowledge, we did not explicitly specify sequences and structures, allowing the network flexibility to learn filters in an unbiased manner. Furthermore, our model explicitly quantifies the strength (in network-defined arbitrary units) of each activated filter to the inclusion or skipping decision. Importantly, we allowed the strength of any filter to vary along the length of an exon, providing the network the flexibility to capture position-dependent effects of RNA features on splicing outcomes.

To arrive at its output, the network computes the difference in the sum total of exon inclusion strengths and exon skipping strengths (Δ strength), which is then converted to predicted PSI. The greater the magnitude of this difference, the closer the PSI is to 0 (difference ≪ 0) or 1 (difference ≫ 0). This additive combinatorial behavior is consistent with previous literature^8,21^.

### Model extends understanding of splicing regulatory logic

Even though our model was trained on a synthetic dataset, it recapitulates and extends domain knowledge from previous genomic and biochemical studies.

Many filters in the model match binding motifs of RNA binding proteins implicated in splicing regulation (splicing factors)^24,25^ (Fig. 2a). Consistent with previous studies, network inclusion filters match binding sites for SR proteins known to promote exon inclusion^23,26^, whereas network skipping filters match binding sites for hnRNP proteins known to promote exon skipping^27^.

**Figure 2.**
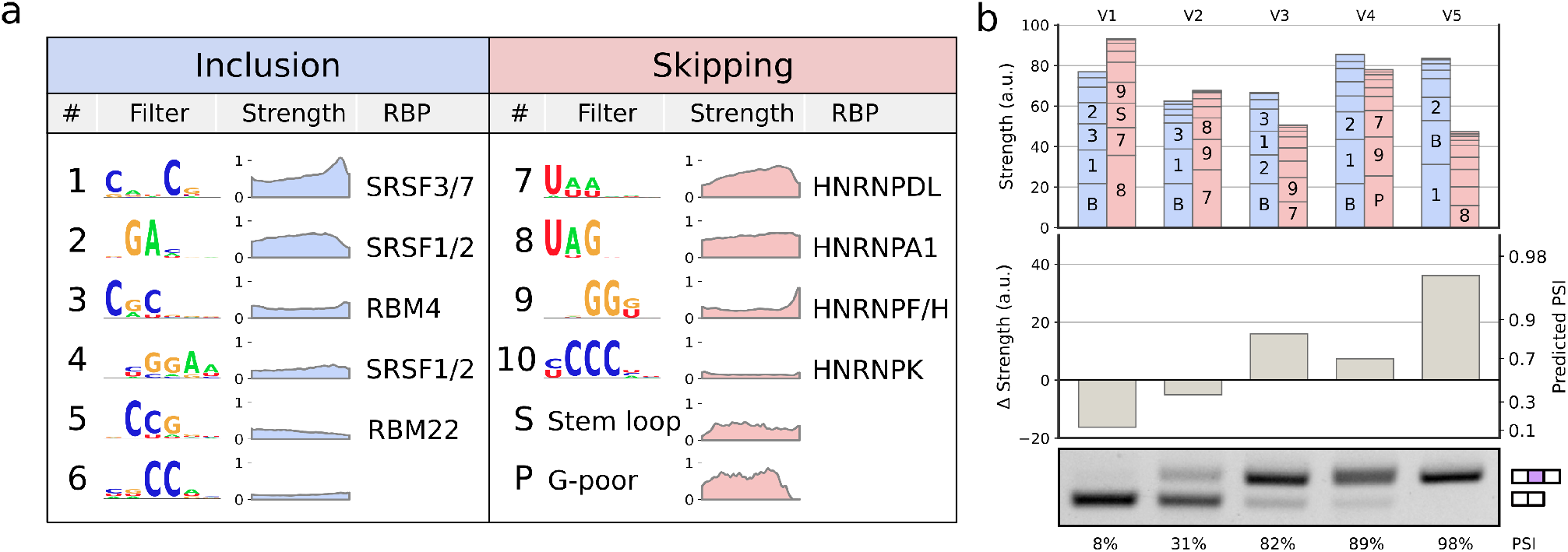
Model expands on known splicing logic and its predictions can be interpreted using balance plots. **a,** Splicing features detected by the model’s filters, represented by their sequence logo^22^. Filters either contribute to inclusion (blue) or skipping (red). Plots show the average strength in our dataset of each filter as a function of position along the exon. RNA binding proteins (RBP) with a similar binding motif, as reported in previous work^23–25^. The model also identified short stem loops and long G-poor stretches as contributing to exon skipping. **b,** Balance plots used to visualize the logic leading to PSI prediction for five randomly picked exons (V1-V5). Bar plot showing the total strength contributed by each filter (top). Bars are labeled by filter numbers from panel **a**. Bar labeled B represents a constant initial inclusion strength. Labels are not shown for smaller bars. The difference between total inclusion and total skipping strengths (Δ strength) leads to predicted PSI (center). PSI as measured by semi-quantitative RT-PCR matches the machine learning predictions (bottom).

However, while the directionality of these RNA features towards splicing was established, their magnitude was not clear. Importantly, the model addresses this issue by assigning a quantitative strength to each filter. Moreover, some filters exhibit striking position dependent strengths, suggesting that the position of an RNA feature along an exon affects its strength.

Surprisingly, our network accurately predicted splicing outcomes using a concise list of filters (Fig. 2a). This contrasts with previous studies suggesting that splicing outcomes result from the combinatorics of hundreds of unique RNA features^8,28,29^.

Using the local interpretability of our model, we introduce a visualization (balance plot) that enables explicit examination and quantification of how multiple RNA features lead to splicing outcomes for any given exon from our dataset (Fig. 2b, Extended Data Fig. 3). For a given exon, the total strengths of activated filters are represented as bars of the appropriate height. Total inclusion strength (blue) and skipping strength (red) are then visible as the height of the stacked bars. The Δ strength is represented by the difference in heights between the stacked inclusion and skipping filters. These visualizations provide an intuitive tool to understand the contributions of individual sequence and structure features leading to each exon’s predicted PSI. They emphasize that splicing logic results from contributions of many RNA features along the exon, and that a single nucleotide can be part of multiple overlapping filters^6,8^.

### Discovery and validation of novel splicing features

Next, we asked whether our interpretable-by-design model could advance scientific discovery by identifying novel splicing features. While most network filters were consistent with previously-described splicing features, two uncharacterized long skipping filters with strong influence on splicing predictions stood out (Fig. 2a). We confirmed that these filters were robustly identified across multiple initialization seeds and training/testing splits, suggesting that they are not training artifacts. We then turned our attention to characterizing and experimentally validating these features.

Examining the first uncharacterized filter revealed that it identifies stem loop structures with short, GC-rich, 5-7 nucleotide double-stranded regions (Fig. 3). Next, we experimentally validated that these stem loops contribute to exon skipping and are not artifacts. We introduced mutations that disrupt double-stranded base pairing in an exon with such a stem loop. First, we introduced single nucleotide mutations predicted to abolish the stem by disrupting base pairing. Notably, these mutations were designed to minimize disruptions of other filters, ensuring that prediction differences are mainly due to altered secondary structure, and not due to the introduction or disruption of other sequence features. In addition to two such mutations, we also introduced both compensatory mutations together, restoring the original stem loop structure^30^. We measured splicing outcomes for all four individual reporters (original, upstream mutation, downstream mutation, and double mutations) and observed that splicing outcomes matched our predictions (Fig. 3). Namely, PSI increased dramatically in both single nucleotide mutants, in agreement with the predicted decrease in filter strength. Furthermore, when both compensatory mutations are present and structure is restored, measured PSI was comparable to that of the original exon. We applied the same experimental validation scheme to two other stem loop-containing exons. In both cases, stem-disrupting single mutations increased exon inclusion and structure-restoring double compensatory mutations had minimal effects (Extended Data Fig. 4). Together, these experiments demonstrate that model-identified stem loops, rather than sequence, contribute to exon skipping.

**Figure 3.**
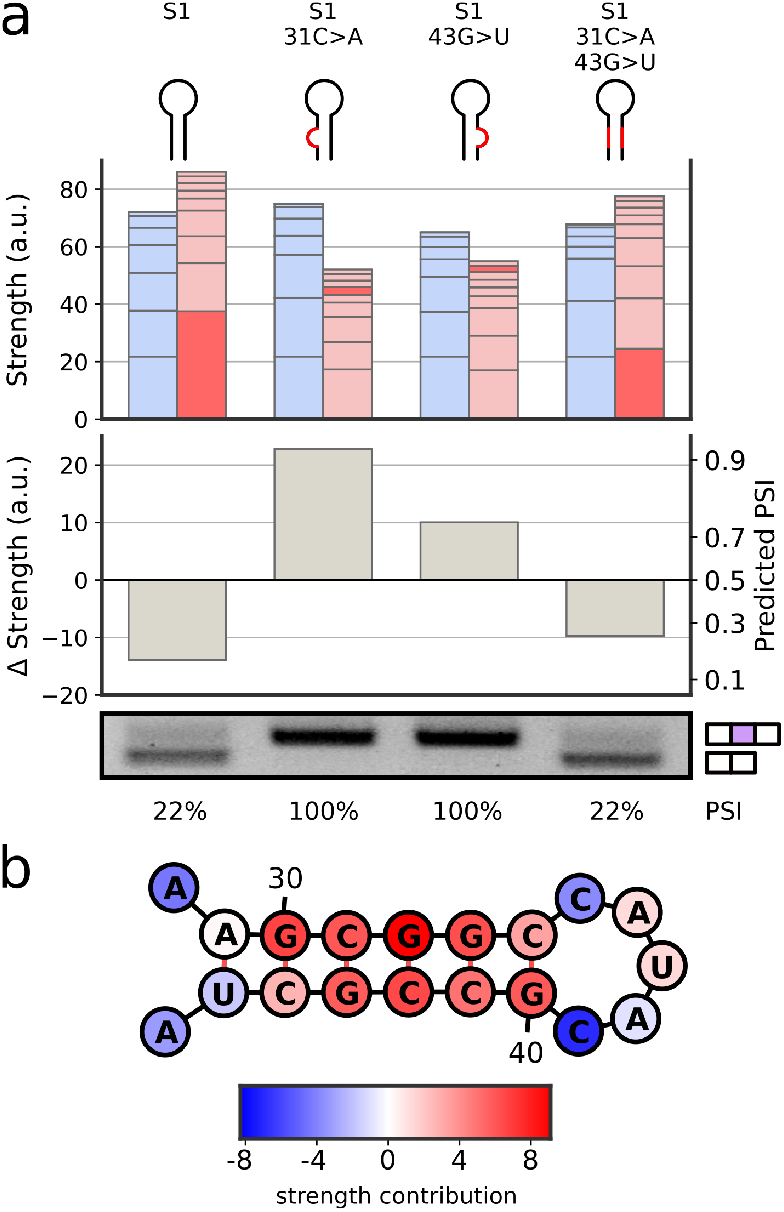
Validation of novel stem loop feature. **a,** The machine learning model identifies a stem loop in an exon (S1) as having a strong skipping strength (dark red bar; top), leading to near complete skipping prediction (middle). Single nucleotide mutations disrupting a downstream or upstream stem base pair are predicted to significantly reduce the skipping strength and restore exon inclusion. Finally, including both single nucleotide mutations is predicted to restore the stem loop skipping strength and lead to skipping. RT-PCR validation (bottom) confirms the machine learning predictions. **b,** The stem loop identified in S1, with the individual contributions to its strength by each nucleotide.

In contrast, examining the second uncharacterized filter did not reveal any secondary structure preference. Instead, the filter exhibited a preference for long guanine depleted (G-poor) sequences (Fig. 4a). To validate that guanine depletion underlies filter behavior, we selected an exon with a G-poor sequence and introduced a single C>G mutation. As before, we ensured that the predicted strengths of other filters are only minimally disrupted (Fig. 4b). Strikingly, this single mutation led to marked increase in PSI. We applied the same validation scheme to three other exons with G-poor sequences; in every instance, a single C>G mutation increased exon inclusion (Extended Data Fig. 5). To the best of our knowledge, a long G-poor sequence has not been described in the literature.

**Figure 4.**
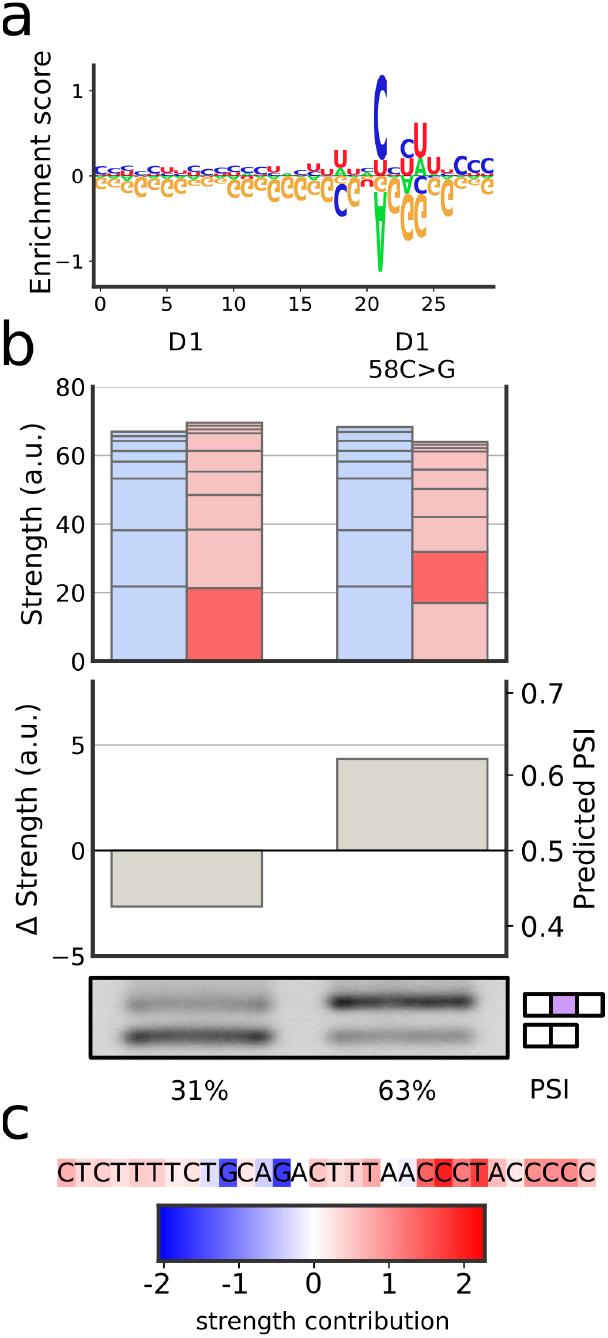
Validation of the G-poor feature. **a,** The G-poor filter, represented by its enrichment-depletion logo^31^. **b,** The machine learning model identifies a G-poor stretch in an exon (D1) as having a strong skipping strength (dark red bar, top), leading to skipping prediction (middle). A single nucleotide C>G mutation is predicted to disrupt the G-poor stretch and restore exon inclusion (right bars). RT-PCR validation (bottom) confirms the machine learning predictions. **c,** The G-poor stretch identified in D1, with the individual contributions to its strength by each nucleotide.

Collectively, these experiments confirm that stem loops and G-poor sequences identified by the model reflect bona fide splicing features.

## Discussion

In this study, we demonstrate that an interpretable-by-design model advanced scientific discovery. Our model accurately predicts splicing outcomes on both our assay and on previously published assays, demonstrating that interpretability need not come at the expense of accuracy or generalizability. Model interpretability enabled a systematic understanding of RNA splicing logic, including the identification of two candidate novel exon skipping features which were subsequently experimentally validated. The model’s ability to quantify contributions of specific features to splicing outcomes for individual exons has considerable potential for a range of medical and biotechnology applications, including genome- or RNA-editing of target exons to correct splicing behavior or guiding rational design of RNA-based therapeutics like antisense oligonucleotides^32^.

In addition, model-identified features hint at novel biochemical mechanisms that warrant further study. For example, the fact that splicing decisions are modeled well by an additive quantity (Δ strength) supports a biochemical mechanism involving the nuclear spatial organization of SR and hnRNP proteins^33^. Furthermore, the novel skipping-promoting G-poor feature may point to an uncharacterized RNA binding protein or complex. These open questions further underscore how interpretable-by-design models can advance scientific discovery by aiding hypothesis generation.

Our model performs well on synthetic datasets from immortalized cell lines, yet further work is needed to capture the dynamics of developmentally regulated splicing logic^34–36^. Importantly, splicing outcomes change depending on the expression level of cell type-specific RNA binding proteins^37^. These questions can be addressed by generation of additional synthetic splicing datasets in developmentally relevant cell types paired with interpretable-by-design models that capture cell type-specific regulatory features.

Beyond the context of splicing, the interpretable-by-design framework can be used to decipher the multiple, complex, and overlapping codes that dictate biomolecular processing. Importantly, many rich synthetic datasets that address RNA untranslated 5‘^38^ and 3‘^39^ region regulation, methylation^40^, and small RNA biogenesis^41^, have already been generated. We expect that additional data generation efforts paired with the interpretable-by-design framework will stimulate advances in understanding biological codes more broadly.

## Data availability

Sequence data that support the findings of this study have been deposited in the NCBI’s Gene Expression Omnibus under accession number GSE200096.

## Code availability

Custom code, preprocessed datasets, and trained model are available on GitHub (https://github.com/regev-lab/interpretable-splicing-model).

## Acknowledgements

We thank members of the Regev laboratory and Lawrence Chasin for feedback; and Megan S. Hogan and Matthew T. Maurano (NYU Institute for Systems Genetics) for technical assistance with amplicon sequencing. We thank Georg Seelig, Jef Boeke, and Brenton Graveley for plasmids used to construct the reporter assay. This work was partly supported by a PhRMA Fellowship (M.S.), Lalor Foundation Fellowship (S.E.L), Life Sciences Research Foundation Fellowship from Additional Ventures (S.E.L.), Simons Investigator Award, and NSF MCB-2226731 (O.R.), and NYU IT High Performance Computing resources, services, and staff expertise.

## Author contributions

The study was initiated by S.E.L. and O.R. Experiments and machine learning analysis were performed by S.E.L. and M.S. All authors wrote, reviewed, and provided feedback on the manuscript.

## Competing interests

The authors declare no competing interests.

## I. Methods

### I.1 Reporter assay design and cloning

The splicing reporter is based on a three-exon beta globin minigene^42^ under the control of a truncated mammalian CAG promoter. The massively parallel splicing assay allows for high-throughput characterization of exon variants on splicing outcomes^43^ using Gibson assembly and ligation cloning. The assay replaces the middle beta globin exon with 70nt random sequences flanked by weak splice sites (MaxEnt scores^44^: 3’ss 9.41, 5’ss 5.06). Each 70nt exon is coupled with a 20nt barcode downstream of the third exon, allowing for identification of middle exon identity in exon skipping products. Briefly, two pools of oligonucleotides (random exons and barcodes with complementary linker regions and flanking overlap sequences for Gibson assembly) were synthesized as single-stranded oligonucleotides (IDT DNA Technologies) and joined using an anneal-extend procedure. 200nM of each oligonucleotide were joined in a 100μL reaction (Phusion® Hot Start Flex 2X Master Mix, NEB). Oligonucleotides were denatured at 98°C for 10min, cooled slowly to 60°C (0.1°C/sec), annealed at 60°C for 5min, and extended at 72°C for 60min. Single-stranded products were removed from pooled double-stranded exon-barcode using silica column purification according to the manufacturer’s specifications (ZymoPURE Plasmid Miniprep Kit). Pooled exon-barcode products were cloned into a backbone digested with BsmBI and XbaI and expanded using electrocompetent bacterial cells (ElectroMAX™ DH10B Cells, ThermoFisher) on large solid agar Bioassay plates (Nunc™ Square BioAssay Dishes, ThermoFisher). After resuspending pooled bacteria in 1X PBS, DNA was recovered using silica column purification (ZymoPURE II Plasmid Maxiprep Kit, Zymo Research) following manufacturer’s specifications. The resulting pooled library (Lib1) includes the truncated CAG promoter, followed by the first minigene exon and intron, and the exon-barcode insertion. High-throughput amplicon sequencing of Lib1 was used to match exon-barcodes pairs. To generate the final splicing reporter assay (Lib2), a fixed sequence, containing the second intron and third exon, was introduced to separate exons from their barcodes. Lib1 was digested with Esp3I (NEB) to introduce overhangs between the exons and barcodes; the digested product was gel-purified to facilitate downstream cloning (Zymoclean Gel DNA Recovery Kits). A segment containing the second intron and third exon was ligated into the digested Lib1 product (NEB Quick Ligation). Lib2 library was expanded using electrocompetent bacteria cells resulting in about 10 times as many colonies as Lib1 to ensure even representation across reporters and recovered using silica column purification as described for Lib1. DNA was quantified using a spectrophotometer (NanoDrop™ One^C^, Fisher Scientific).

### I.2 Individual reporter cloning

To validate consequences of point mutations on splicing outcomes, individual exons were synthesized as two single-stranded oligonucleotides (IDT DNA Technologies) and joined using an anneal-extend procedure. Briefly, 200nM of each oligonucleotide were joined in a 100μL reaction with 5U DNA polymerase (NEB Klenow). Oligonucleotides were denatured at 98°C for 10min, annealed after cooling slowly to 37°C (1°C/sec), and extended at 37°C for at least 2 hours. Reactions were heat inactivated at 75°C for 20min and used directly for Gibson assembly into a digested receiving plasmid with a fixed barcode.

### I.3 Cell culture

HeLa cells (ATCC) were grown in high-glucose DMEM medium supplemented with 10% fetal bovine serum and penicillin and streptomycin (ThermoFisher). All cells were grown at 37°C, 5% CO2, and 95% relative humidity.

### I.4 Transfection, RNA extraction, and reverse transcription

Cells were transfected at 60-80% confluence with FuGENE HD® according to the manufacturer’s protocol at a 3:1 FuGENE HD® to DNA ratio. For high-throughput measurements of splicing outcomes, 10 μg pooled reporter assay DNA was transfected in three 100 mm plates. For biochemical analysis of individual reporters, 1 μg or 2.5 μg individual reporter DNA was transfected into each well of a 12- or 6-well plate (respectively). 24 hours after transfection, total RNA was isolated from detached cells (Accutase^®^, ThermoFisher). For amplicon sequencing, total RNA was isolated using phenol-chloroform (Ambion) extraction (5PRIME Phase Lock Gel, Quantabio) followed by DNase treatment (TURBO DNase). For biochemical analysis, RNA was isolated using a silica column (illustra™ RNAspin Mini RNA Isolation Kit, GE Healthcare) with on-column DNase digestion following manufacturer’s automated protocol. DNase-treated RNA was reverse transcribed using a reporter-specific primer following manufacturer’s specifications (SuperScript IV Reverse Transcriptase, Thermo Fisher) with RNase H treatment. Reverse transcription primers included degenerate nucleotides to serve as unique molecular identifiers (UMIs) during amplicon sequencing^45,46^. cDNA products were used for amplicon sequencing or biochemical analysis.

### I.5 Amplicon sequencing

Amplicon sequencing was used to identify exon-barcode pairings in Lib1 and to quantify splicing products from reverse-transcribed cDNA. Second-strand synthesis added additional UMIs in a single anneal-extend cycle of 98°C for 10min, cooled slowly to 60°C (0.1°C/sec), annealed at 60°C for 5min, and extended at 72°C for 5min (Phusion® Hot Start Flex 2X Master Mix, NEB). Resulting double-stranded amplicons were amplified using a two-stage procedure. In the first stage, targets were amplified by PCR primers. PCR was performed using the following protocol: 98°C for 30sec initial denaturation, then 16 cycles of 98°C denaturation for 10sec, 60°C annealing for 15sec, 72°C extension for 1min 45sec, and a final extension step at 72°C for 5min (Phusion® Hot Start Flex 2X Master Mix, NEB). Longer extension times and minimal number of PCR cycles were used to avoid recombination across exons and barcodes. The number of cycles was determined for each sample by first running 10μL qPCR reactions (LightCycler® 480 SYBR Green I Master, Roche). In the second stage, index primers were added using 5 PCR cycles. PCR was performed using the following protocol: 98°C for 30 s initial denaturation, then 5 cycles of 98°C denaturation for 10sec, 71°C annealing for 15sec, 72°C extension for 1min 45sec, and a final extension step at 72°C for 5min (Phusion^®^ Hot Start Flex 2X Master Mix, NEB). Final DNA concentrations were measured using fluorometric measurements (Qubit 1X dsDNA HS Assay, Thermo Fisher) on a Qubit 3 fluoremeter. Paired-end sequencing was carried out on an Illumina NextSeq 550 with 10% PhiX spiked in, with 54 cycles in read 1 (reverse) and 106 in read 2 (forward). About 4M paired-end reads (> 10X coverage) were acquired for Lib1 exon-barcode sequencing and an average of 22M paired-end reads (> 50X coverage) for each PSI quantification replicate.

### I.6 Biochemical analysis

PCR amplification reactions to determine splicing products were carried out in 20μL reactions containing 10μL OneTaq® 2X Master Mix with Standard Buffer (NEB), 200 nM each forward and reverse primers (IDT), and 1μL cDNA. PCR was performed using the following protocol: 94°C for 30 s initial denaturation, then 25 cycles of 94°C denaturation for 10 s, 62°C annealing for 15 s, 68°C extension for 20 s, and a final extension step at 68°C for 1 min. 5μL final PCR product was run out on 2.0% agarose (Denville Scientific) Tris-acetate-EDTA (TAE) gel with ethidium bromide and visualized on a Bio-Rad imager. Densitometry measurements to calculate PSI were measured using Bio-Rad Image Lab (Windows v6.1).

### I.7 Reporter assay preprocessing

The list of all exons in the reporter assay with their corresponding barcodes was extracted from DNA sequencing of Lib1. To ensure unique coupling of barcodes to exons, barcodes appearing with more than one exon sequence were filtered out. This step ignored exon sequences appearing only once, as those are likely due to sequencing errors. Barcodes with fewer than two DNA reads in total were also filtered out.

Next, splicing outcomes were extracted from RNA sequencing of each of the three replicate transfections of Lib2. For each replicate, each read was identified by barcode and was assigned a splicing outcome (exon skipping, exon inclusion, intron retention, splicing inside exon, or unknown splicing). Carryover from Lib1 was filtered out, as were reads for which exon 1 could not be identified. Using unique molecular identifiers (UMIs), the fraction of duplicate reads in each replicate was estimated to be below 23%. The counts from all three replicates were merged for downstream analysis. Barcodes with fewer than 60 total reads, barcodes that contained an Esp3I restriction site in either strand of the exon or its barcode, and barcodes where inclusion and skipping made up less than 80% of all reads were filtered out.

Finally, the dataset was generated by computing PSI for each barcode as

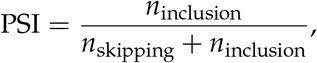

where *n*_inclusion_ is the total number of exon inclusion reads, and *n*_skipping_ is the total number of exon skipping reads. In addition to the measured PSI, the dataset includes for each barcode: (1) a 90 nucleotide sequence, containing the 70 nucleotide variable exon sequence plus the 10 fixed flanking nucleotides on each side; (2) structure in dot-bracket notation predicted by RNAFold (Vienna RNA^20^, version 2.4.17), using default parameters; (3) an indicator vector indicating which nucleotide participates in a predicted G-U wobble base pair. The dataset was split randomly into a training set and a test set in an 80/20 split, using a fixed seed for reproducibility.

### I.8 Model design

The model’s input is a triple of vectors (*x*_seq_, *x*_struct_, *x*_wobble_),

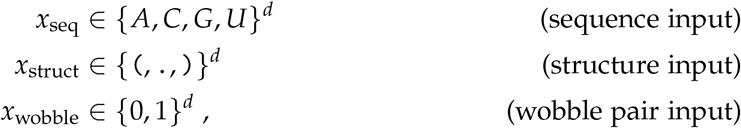

where *d* = 90. The neural network contains four “strength-computation modules” (SCM) defined as

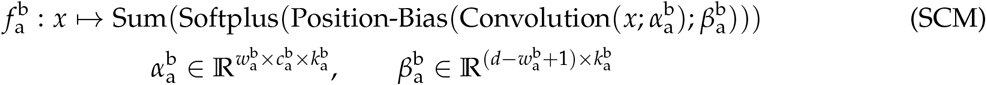

where a ∈ {incl, skip}, and b ∈ {seq, struct}. The input is either *x* = [*x*_seq_] (sequence SCM) or *x* = [*x*_seq_, *x*_struct_, *x*_wobble_] (structure SCM). The 1D convolutional layer contains 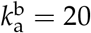 convolutional filters of width 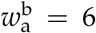 for each sequence SCM (b = seq), and 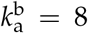 convolutional filters of width 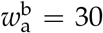 for each structure SCM (b = struct). The number of input channels is 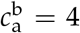 for sequence SCM (corresponding to the one-hot encoded four nucleotides) and 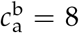 for structure SCM (corresponding to sequence, structure, and wobble indicator). The output of the convolution layer is a 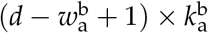 matrix *z* of “raw” strengths. The position bias layer maps inputs *z* to 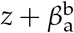, adjusting the raw strengths based on position along the exon. Finally, each position-adjusted raw strength is passed through a softplus activation, and the resulting strengths are all summed up to form the output of the SCM 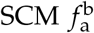.

The splicing prediction model *m*(*x*_seq_, *x*_struct_, *x*_wobble_; *θ*) is then defined as

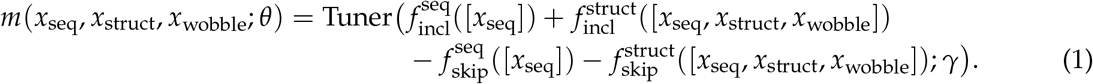

This model computes the total strength for inclusion and for skipping and uses their difference to predict splicing outcomes. The function Tuner(·; *γ*) : ℝ → [0,1] is a learned nonlinear activation function that maps this difference to a PSI prediction. It consists of a 3-layer fully connected network with a residual connection from the input to the output layer, followed by a sigmoid activation. The parameter set *θ* contains the parameters of all SCMs and the parameter *γ*.

### I.9 Model training

The model was implemented in Python 3.8^47^ using Tensorflow 2.6^48^ and Numpy 1.20^49^. Batched gradient descent was used to optimize the model’s parameters using the Adam optimizer, with KL divergence as the loss function. Hyperparameters such as regularization parameters were tuned with grid search. Training the model took about 2 hours on a mid-range 4-core with 16GB of RAM.

To improve interpretability, the model was trained in steps (custom training schedule), progressively adding learnable parameters in each step. First, a simplified model given by

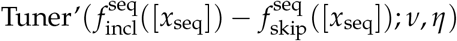

was trained. Here, Tuner’(·; *v, η*) : ℝ → [0,1] is a learned nonlinear activation function defined by *x* → *σ*(*vx* + *η*) where *σ* is the sigmoid function, and *v* and *η* are two real parameters. This step ensures that short sequence motifs are captured by the sequence SCMs and not the more complex structure SCMs. In the second step, the structure SCMs were added, leading to a model identical to the final one (1), except for the use of Tuner’ instead of Tuner. The sequence SCM weights were initialized to those of the previous model. In the third and last step, the Tuner function was introduced, leading to the final model (1). SCM weights were initialized to those of the previous model.

To further improve the model’s interpretability, regularization terms were added. First, to obtain a concise list of filters, an activity regularization loss term was used. The term consists of the ℓ_1_ norm of all the strengths. Second, a smoothness regularization loss term was applied to position bias layer weights. This term consists of the ℓ_2_ norm of the discrete derivative of the weight vectors (defined as the difference between the vector and itself shifted by one along the sequence dimension). Each of the two loss terms was multiplied by a hyperparameter.

Hyperparameters were optimized based on two criteria: held-out KL divergence and sparsity of activations. Sparsity was measured as the minimum number of activations needed per exon to achieve KL below a threshold. Among all hyperparameters leading to sufficiently high accuracy and sparsity, the one with the highest smoothness regularization was chosen.

### I.10 Prediction accuracy on other assays

Exon sequences and PSI measurements were obtained from previous publications. Exons including indel mutations or differing from WT sequence in the first or last three nucleotides were filtered out. Sequences were padded to 70 nucleotides, and predicted PSI was then computed using our model. To account for differences in splice sites, flanking sequences, and cell types, one correction term was introduced per assay, as described previously^16^.

### I.11 Filter visualization

To avoid reporting redundant sequence filters, hierarchical clustering using Scipy^50^ was applied. Each sequence filter was represented by a vector containing its total strength for each of the exons in the dataset. The strongest filter in each cluster was then used to generate a sequence logo^22^. The logo represents the set of 6-mers that lead to positive filter activation.

The structure filters included one G-poor filter and three stem loop filters. Since enumerating all 30-mers is not tractable, the G-poor sequence logo was computed by evaluating the filter on a subset of sequences from our dataset. As the three stem loop filters differed in the length of the loop (short, medium, long) but were otherwise very similar, they were considered as one cluster. Layer-wise relevance propagation was used to visualize individual nucleotide contributions to filter strength^51^.

### I.12 Ruling out sequencing artifacts

The reduction in measured PSI associated with the presence of the stem loop and G-poor stretch features could potentially be a technical artifact due to decreased amplification or sequencing efficiency of exon inclusion products. To rule this out, we verified that the presence of these features was not accompanied by a decrease in the total number of sequencing reads, and was instead accompanied by an increase in the number of exon skipping reads (Extended Data Fig. 6).

### I.13 Design of mutant constructs

To validate the stem loop feature, candidate exons with high medium-length stem loop filter strength (top percentile) but with no other stem loop activations elsewhere in the exon were selected. Three mutants of each such exon were then generated. To ensure these mutants do not introduce or disrupt other features, exons where this mutation significantly changed strengths of other filters were filtered out.

To validate the G-poor stretch feature, candidate exons that strongly activate the G-poor filter exactly once along the exon were selected. For each candidate exon, a C-to-G mutation in the middle of the activated filter’s window was introduced. As before, to ensure this does not introduce or disrupt other features, exons where this mutation significantly changed strengths of other filters were filtered out.

## II. Extended Data Figures

**Extended Data Figure 1.**
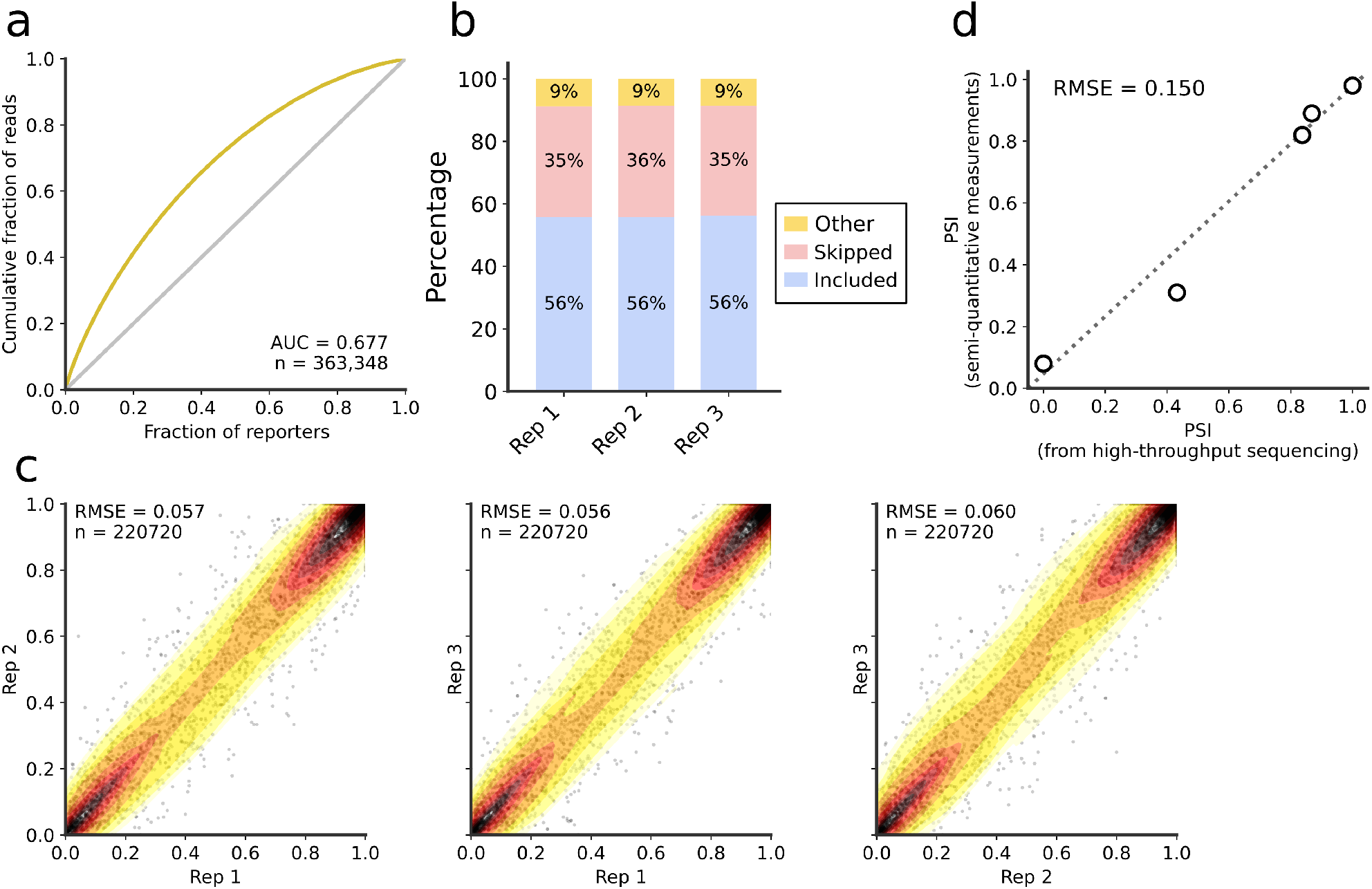
Assay quality checks. **a,** Lorenz plot showing the distribution of reads across the reporters in a high-throughput sequencing of the library DNA (gold). A perfectly even library would be on the diagonal (gray). **b,** Over 90% of splicing products corresponded to exon inclusion or exon skipping products, as measured by high-throughput RNA sequencing. **c,** Comparison of PSI measurements across the three biological replicates. **d,** Comparison of PSI measurements from high-throughput sequencing and semi-quantitative measurements for five individual reporters (V1-V5). AUC: area under curve; RMSE: root-mean-square error.

**Extended Data Figure 2.**
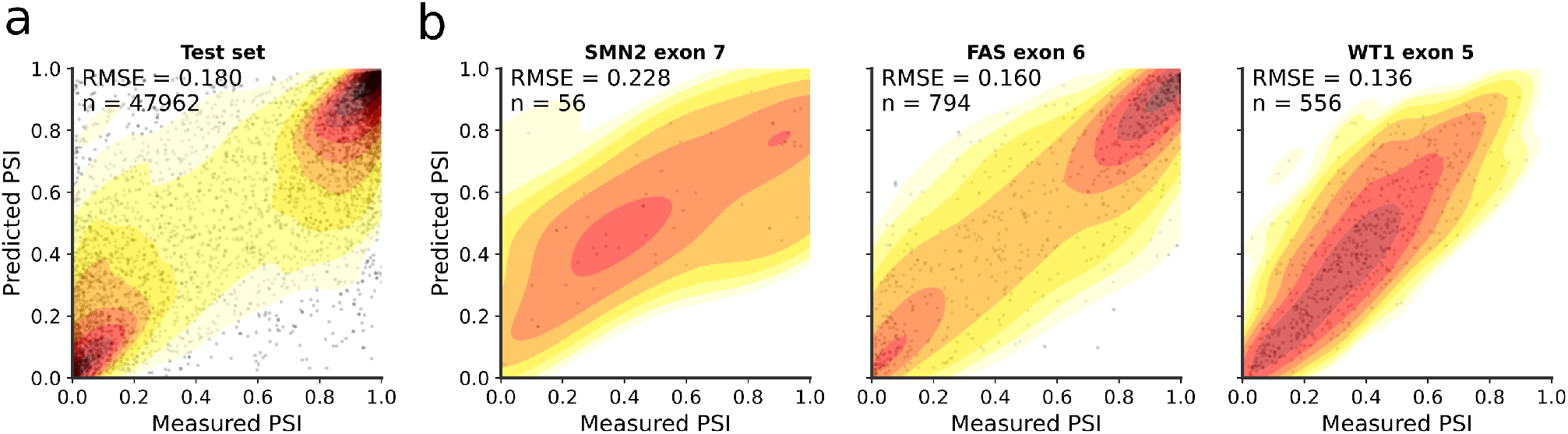
Model predictive accuracy. **a,** Predictions on the held-out experimental data. **b,** Predictions on previously-published assays: SMN2 exon 7 (C33a cells)^8,13–15^, FAS exon 6 (HEK293 cells)^16^, and WT1 exon 5 (HEK293 cells)^7^.

**Extended Data Figure 3.**
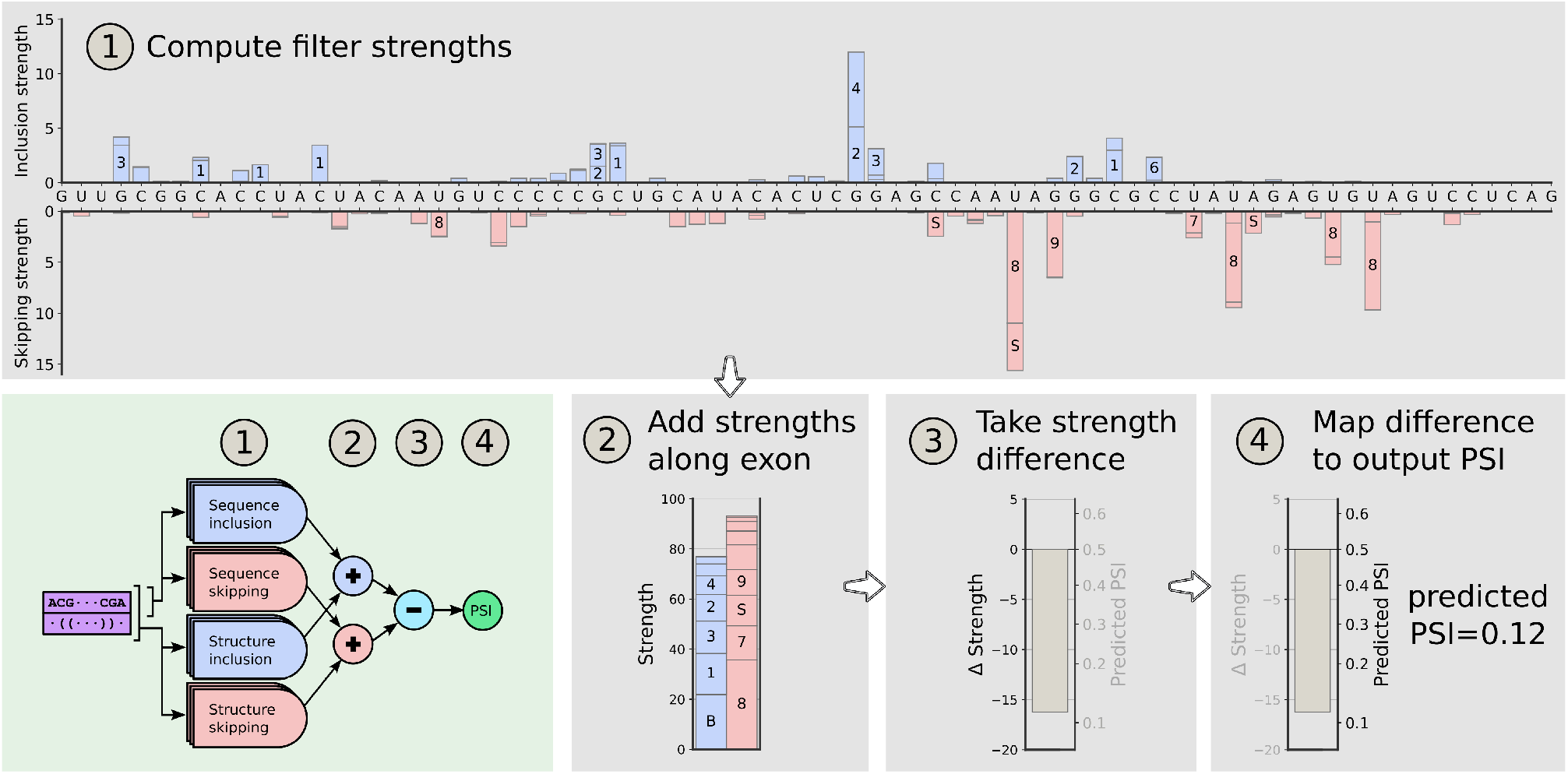
Tracing the neural network computation from exon sequence to predicted PSI. Our neural network (bottom left) arrives at its predictions in four steps. First, using exon sequence and predicted structure, it computes filter strengths for each position along the exon (1). Next, it adds all inclusion strengths together and all skipping strengths together (2). Then, the difference between these two strengths (Δ strength) is computed (3). Finally, it maps that difference to a predicted output PSI (4).

**Extended Data Figure 4.**
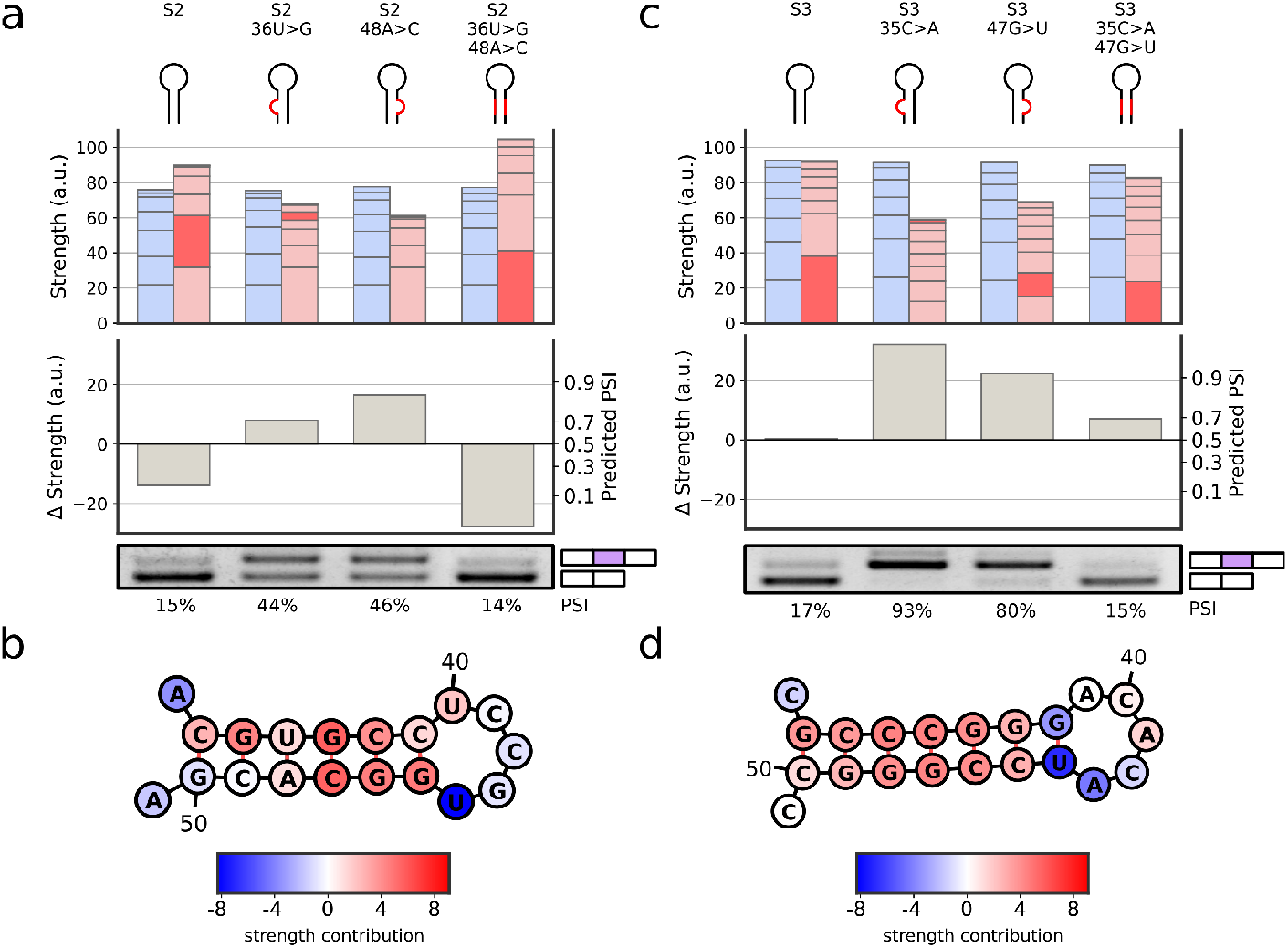
Additional validation of novel stem loop feature. Two more exons (S2, S3) were chosen and validated as in Figure 3.

**Extended Data Figure 5.**
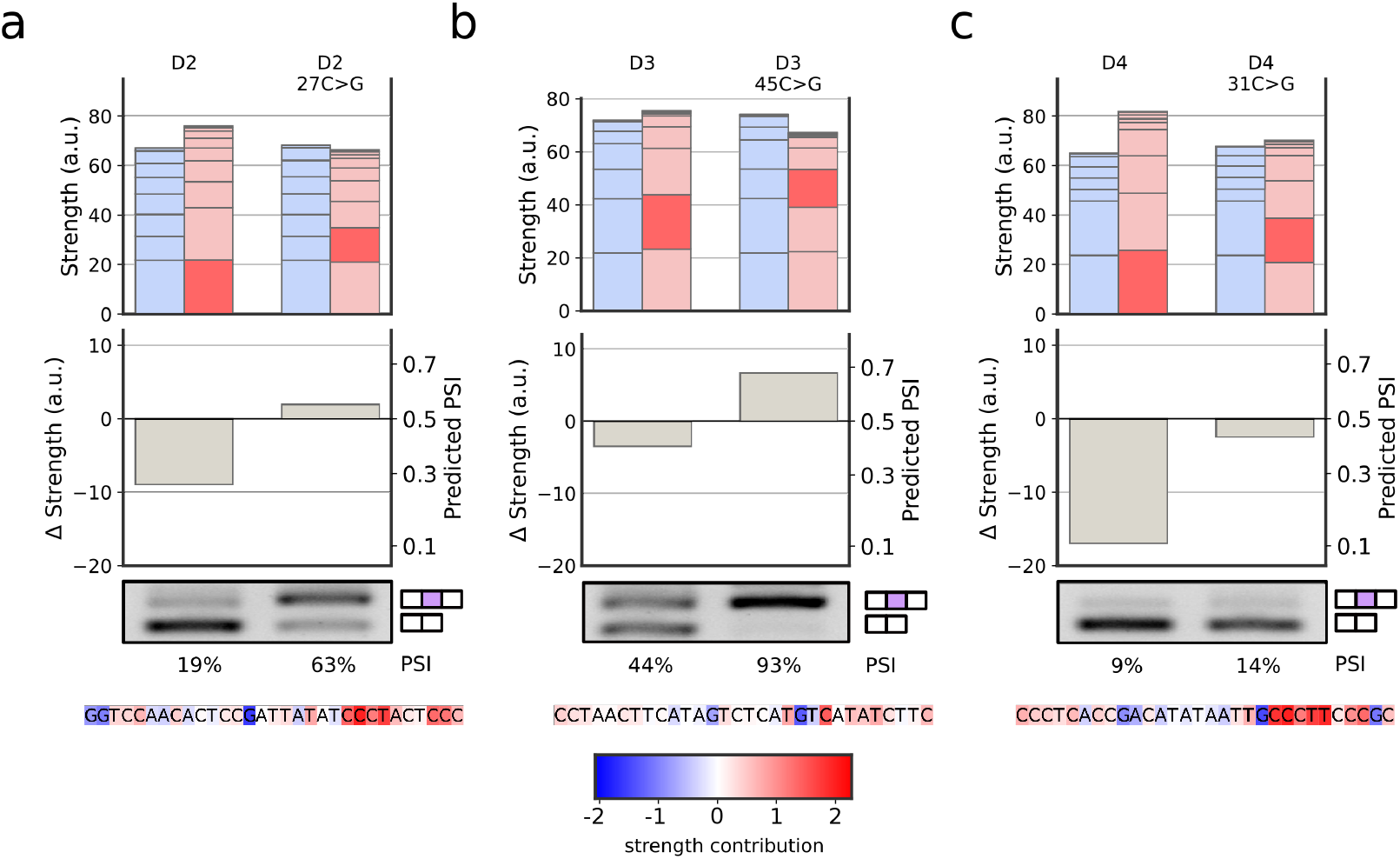
Additional validations of the G-poor feature. Three more exons (D2, D3, D4) were chosen and validated as in Figure 4.

**Extended Data Figure 6.**
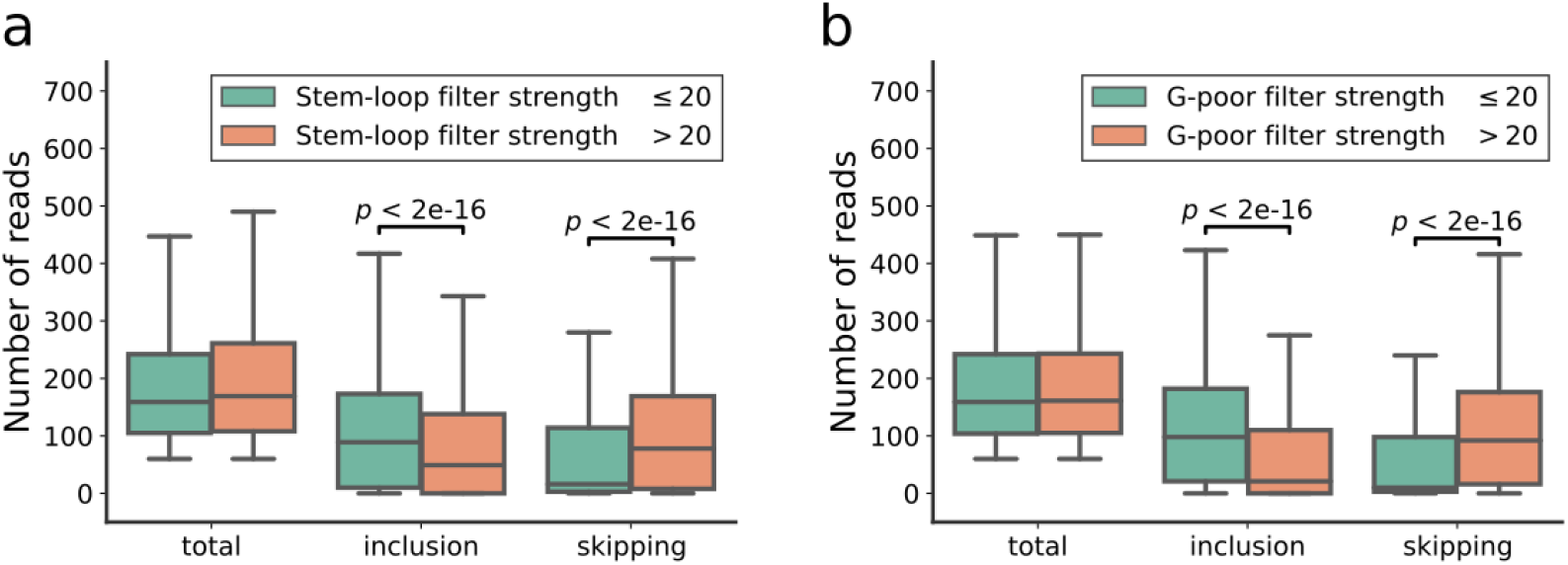
Effect of novel features on absolute read counts. **a,** Box plot showing the distribution of the total number of sequencing reads, the number of exon inclusion reads, and the number of exon skipping reads, for exons with stem loop strength at most 20 and greater than 20. **b,** As in panel **a** for G-poor strengths. Center line: median; box limits: upper and lower quartiles; whiskers: 1.5x interquartile range. *p* values: Student’s t-test.

## Notes

https://www.ncbi.nlm.nih.gov/geo/query/acc.cgi?acc=GSE200096

https://github.com/regev-lab/interpretable-splicing-model

### Competing Interest Statement

The authors have declared no competing interest.

